# Comparing physiological impacts of positive pressure ventilation versus self-breathing via a versatile cardiopulmonary model incorporating a novel alveoli opening mechanism

**DOI:** 10.1101/2024.03.12.584596

**Authors:** MT Cabeleira, DV Anand, S Ray, C Black, NC Ovenden, V Diaz-Zuccarini

**Affiliations:** Department of Mechanical Engineering, University College London, London, WC1E 7JE, UK; Department of Mathematics, University College London, London WC1E 6BT, UK; Paediatric Intensive Care Unit, Great Ormond Street Hospital for Children NHS Foundation Trust, London, UK; University College London Hospitals NHS Foundation Trust, London NW1 2BU, UK

**Keywords:** Cardiopulmonary model, lung recruitment, positive pressure ventilation, PEEP

## Abstract

Mathematical models can be used to generate high-fidelity simulations of the cardiopulmonary system. Such models, when applied to real patients, can provide valuable insights into underlying physiological processes that are hard for clinicians to observe directly. In this work, we propose a novel modelling strategy capable of generating scenario-specific cardiopulmonary simulations to replicate the vital physiological signals clinicians use to determine the state of a patient. This model is composed of a tree-like pulmonary system that features a novel, non-linear alveoli opening strategy, based on the dynamics of balloon inflation, that interacts with the cardiovascular system via the thorax. A baseline simulation of the model is performed to measure the response of the system during spontaneous breathing which is subsequently compared to the same system under mechanical ventilation. To test the new lung opening mechanics and systematic recruitment of alveolar units, a positive end-expiratory pressure (PEEP) test is performed and its results are then compared to simulations of a deep spontaneous breath. The system displays a marked decrease in tidal volume as PEEP increases, replicating a sigmoidal curve relationship between volume and pressure. At high PEEP, cardiovascular function is shown to be visibly impaired, in contrast to the deep breath test where normal function is maintained.

## 1 INTRODUCTION

Mathematical models and computational simulations are crucial in enhancing our understanding of health, physiology, and therapeutic strategies, playing a pivotal role in biomedical research and clinical applications [1, 2]. A wide variety of physiological models have been developed, ranging from simple zero-dimensional models, containing little if any spatial detail, to advanced three-dimensional multiscale simulations that intricately attempt to capture the vast complexities of certain specific physiological functions. Typically, the physiological system in a zero-dimensional (lumped parameter) model is conceptualized as a series of discrete compartments, with their dynamics characterized by ordinary differential equations (ODEs)[3]. In contrast, one-dimensional (1D), two-dimensional (2D), and three-dimensional (3D) models employ partial differential equations (PDEs), that additionally capture potentially important spatial dependence on certain physiological processes and mechanisms. Recently, there has been a surge in developing digital twins or virtual patients to facilitate personalized care and treatment. Lumped parameter models play a significant role in creating digital twins, enabling seamless integration of multiple organ systems to mimic the most pertinent physiological processes in real-time [4].

Generally, the lumped parameter models exploit the fluid-electrical-circuit analogy, relating network pressures and flows to mathematically represent the physiological system [5]. These models have a broad range of applications, from studies on single blood vessels to simulations tailored for large vascular networks. In recent advancements within cardiovascular (CV) system modeling, there is a trend towards combining 0D models, often based on the Windkessel model, with 1D and 3D simulations [6]. An example of this is the CircAdapt approach, which employs 0D models for the vasculature coupled to a 3D model for the heart chambers, and offers a comprehensive view of cardiac function [7]. To model CV pathologies more accurately, some studies have adapted their systems of equations to exhibit non-linear behaviors in order to include mechanisms such as ventricular interaction, valve dynamics, and the autonomic nervous system [8, 9]. Similar to CV modeling, models of the pulmonary system (PS) also use the lumped parameter approach to effectively represent various regions within the lungs [10]. More recent approaches have been developed to include multi-branching trees of bronchi, offering a more anatomically detailed structure and function of the PS [11, 12]. A critical aspect of these branching models is the individualized Threshold Opening Pressure (TOP) for each alveolar unit. Yuta et al., [13] proposed a model incorporating 60,000 airway and alveolar units, each with its own TOP following a normal distribution. This model was then applied to explore optimal mechanical ventilation strategies for Acute Respiratory Distress Syndrome (ARDS) patients [14, 15].

Owing to the similar modeling principles of the CV and PS, various integrated cardio-pulmonary (CP) models have been proposed. These models are used to understand the interconnected dynamics of these two systems, including the effects of blood gas levels on heart rate and ventilation, and the influence of respiratory mechanics on venous return and arterial pressure. Consequently, these models are pivotal to investigate the physiological impacts of ventilation and can help determine the most effective, patient-specific ventilation strategy. Flumerfelt et al. [16], introduced a two-compartment lung model integrated with capillaries for gas exchange. This idea was expanded upon by Liu et al. [10] and further advanced by Lu et al. [17], with the latter introducing a detailed CP model. This model encompassed a wide range of physiological aspects, including the mechanics of atria and ventricles, the hemodynamics of both systemic and pulmonary circulations, the baroreflex mechanism for arterial pressure regulation, the mechanics of airways and lungs, and gas exchange at the alveolar-capillary interface, thereby providing a comprehensive view of CP interactions. The works of Ursino et al. [18] laid the foundation for another family of CP models, beginning with a CV model featuring autonomic control and later expanded by Magroso et al. [19] into a CP model that explored the role of carbon dioxide in CV regulation. Chbat et al [20] extended this model to include hypercapnic respiratory failure. Subsequently, Albanese et al [21, 22] further refined the model to include the autonomic nervous system interactions. Other notable CP models in the literature, featuring chemoreceptor and baroreceptor regulation, include those by Fernandes et al. [23] and Lin et al. [24].

Although these models offer valuable insights into the effects of ventilation on CP physiology [17, 19, 22, 25], they often fail to adequately represent the coupled physical interactions between the CV and PS. In particular, the impact of positive pressure ventilation (PPV) on the CP system, which is extremely relevant in intensive care, has not been well explored in the current literature. This is despite the numerous existing studies that indicate prolonged or excessive use of PPV can adversely affect the integrity and functionality of the respiratory system[26]. Patients with conditions like ARDS are notoriously difficult to ventilate efficiently [27] and usually patient-specific strategies are needed to ensure optimal ventilation for these patients [28]. A typical characteristic observed in these patients is lung heterogeneity, which causes some regions of the lung to inflate more than others due to fluid accumulation or damaged tissues during mechanical ventilation [29]. This can potentially lead to overdistension in certain areas while leaving others collapsed or underventilated, resulting in impaired gas exchange and reduced lung compliance [30]. While it has been observed that lung heterogeneity reduces under increasing airway pressure, the extent of the underlying structural heterogeneities persists when pressure is reduced [31]. High PEEP can also affect the CV system, where the increased thoracic pressure can compromise the capacity of the heart to maintain a healthy cardiac output CO [32, 33]. To this end, we aim to develop a CP model that accounts for lung heterogeneity and the aforementioned CP interactions, and precisely quantifies the associated changes in the CV system, such as venous return and cardiac outputs, during both spontaneous breathing and mechanical ventilation.

Our focus is to investigate the effects of Positive Pressure Ventilation (PPV) on the CP system and to compare this ventilation approach with Spontaneous Breathing (SB). To achieve this, we develop an efficient computer implementation of the CP model, leveraging object-oriented programming to conceptualise the model’s architecture into a strategic assembly of generic compartments. This methodology enables the construction of diverse physiological model configurations, that can be easily tailored to study various physiological systems. We exemplify this versatility by developing an integrated mathematical model of the human CP system, comprising a tree structure for the PS with multiple independently configurable alveoli, that demonstrates the adaptability and effectiveness of our methodology. We then apply our CP model to investigate a novel alveoli opening mechanism, uncovering the different modes of operation during the PEEP challenge test and deep breath scenario. The paper is structured as follows: In Section 2 we outline the model generalization technique, followed by a detailed explanation of the equations used in the model and their physiological significance. In Section 3 baseline simulations within the normal physiological range for an adult are presented to validate the model. We then present a Positive End-Expiratory Pressure (PEEP) challenge test and describe the alveolar operating mechanism. Further, we compare the PEEP challenge test with a deep breathing scenario induced by the respiratory muscles. These studies demonstrate that our CP model can simulate the effects of the lung heterogeneity in both PPV and SB realistically. To conclude, in Section 4 we discuss both the applicability and limitations of the model framework presented.

## 2 MATERIALS AND METHODS

The model of the CP system developed in this study integrates key aspects of CV circulation, respiratory mechanics, and cardio-pulmonary interactions. This model can reproduce physiological behaviours commonly seen in adult humans, both under spontaneous breathing and mechanical ventilation, while elucidating the underlying respiratory mechanisms and haemodynamics. Our CP model utilizes a compartmental modelling approach, wherein the CP system is segmented into a series of interconnected compartments, each corresponding to specific sections of the heart, lungs, and vascular network. In this approach, each compartment is effectively a standalone module, tailored to capture the unique physiological functions pertinent to its respective anatomical regions. For example, the trachea is conceptualized as a compartment, that allows airflow from the atmosphere into the bronchial pathways. Similarly, the left heart can be seen as an individual compartment, responsible for receiving blood from the pulmonary veins and channeling it into the systemic arteries. Therefore, the compartments constitute the foundational units, or *generic compartments*, in our model, offering a generalised representation of the various elements within the CP system.

A generic compartment, comprising compliances and resistances to simulate the inflow and outflow behaviour of fluid in a compartment, is shown in Figure 1. The underlying governing equations used to compute the pressure *P*, flow *Q*, and volume changes *V −V*_0_ are as follows:

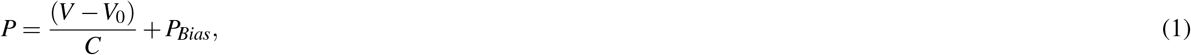

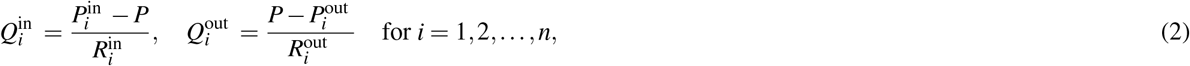

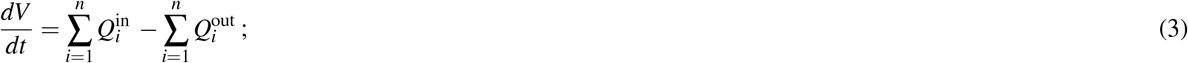

these equations are formulated based upon principles of conservation laws, established constitutive equations, and phenomeno-logical relations. Equation (1) is the constitutive equation that computes the pressure in a compartment based on its compliance and the volume it occupies at that instant of time. The flow in and out of a compartment is then calculated using equation (2), which uses the pressures calculated previously along with resistances that modulate the flow. Equation (3) is the conservation equation that describes the dynamics of air/blood inflow, outflow, and accumulation within the system, maintaining the integrity of volume conservation.

**Figure 1.**
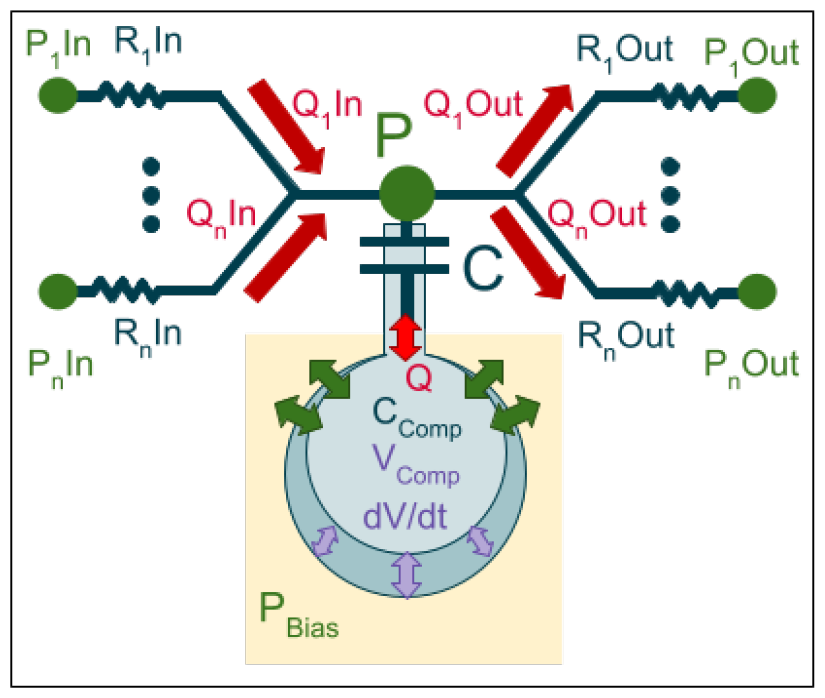
Generic representation of a compartment, composed of a set of resistors that connect the compartment to other compartments and a capacitor that holds a certain volume of fluid. *P* is the pressure felt by the capacitor, *P*^*in*^ and *P*^*out*^ are the pressures of the neighbouring capacitors that are connected to this compartment, *R*^*in*^ and *R*^*out*^ are the resistances of the connecting resistors. *Q*^*in*^ and *Q*^*out*^ represents the inflow and outflow of fluid. C represents the capacity/compliance of the compartment.

It is clear from these three equations that the compliance and resistances dictate the temporal evolution of the pressure and volume in the compartment. The compliances and resistances can be constant (passive) or variable and can also have non-linear relationships with the physiological variables depending on the specific compartment and their interactions with other compartments. Many generic compartments are then strategically assembled to construct a comprehensive model of the CP system. Whilst the temporal dynamics of each compartment are described using ODEs, the spatial dependencies between different compartments are approximated through the strategic implementation of *lumped* properties and the interconnections between the compartments. Despite its simplicity, we demonstrate how this approach permits a detailed exploration of coupled physiological mechanisms, such as respiratory-induced variations in blood flow, and alveolar recruitment mechanisms.

### Cardio-pulmonary system model

An integrated CP model having a lung and cardiovascular system, assembled with appropriate compliances and resistances is shown in Figure 2. To achieve realistic physiological behaviour, the capacitors are categorized into passive, semi-compliant, and variable types, each tailored to represent different compliance characteristics of the CP system. Similarly, the resistances are categorized as either passive or diode-like, enabling the simulation of a broad range of physiological flows and valve dynamics. This diverse array of components ensures that our model not only reflects the complexities of the CP system but also adapts dynamically to simulate realistic physiological mechanisms and processes. Although the assembly of multiple compartments leads to a sophisticated architectural design, it is important to note that each compartment in the CP model is equipped with a capacitor, while resistors establish inter-compartmental connections. This configuration allows the CP model to be conceptualized as a graph, where nodes (capacitors) represent the individual compartments and the resistors are depicted as edges linking these nodes. From a computational perspective, the graph-based architecture facilitates a matrix representation that encodes the connectivity map between different compartments. This methodology effectively simplifies the representation of the model’s architecture, thereby streamlining the construction and manipulation of the interconnected CP system. In our CP model, each compartment is designated to represent a distinct segment of the heart, lungs, and vasculature. The CV system is segmented into nine distinct compartments. These include two compartments for the ventricles (denoted as Hl and Hr), systemic arterial, capillary, and venous compartments (denoted as As, Cs, and Vs respectively), and a thoracic venous compartment (Vt), representing the veins within the thorax. Additionally, it encompasses compartments for the pulmonary arterial, capillary, and venous systems (Ap, Cp, and Vp). The entire pulmonary vasculature, the heart, and the thoracic vein are all housed within the thoracic cavity. The PS is composed of a trachea, bronchi, and multiple independently configurable alveolar compartments enclosed in the pleural space. In the following subsections, we describe the compliance and resistance choices associated with the vasculature, heart, lung, and thoracic cavity.

**Figure 2.**
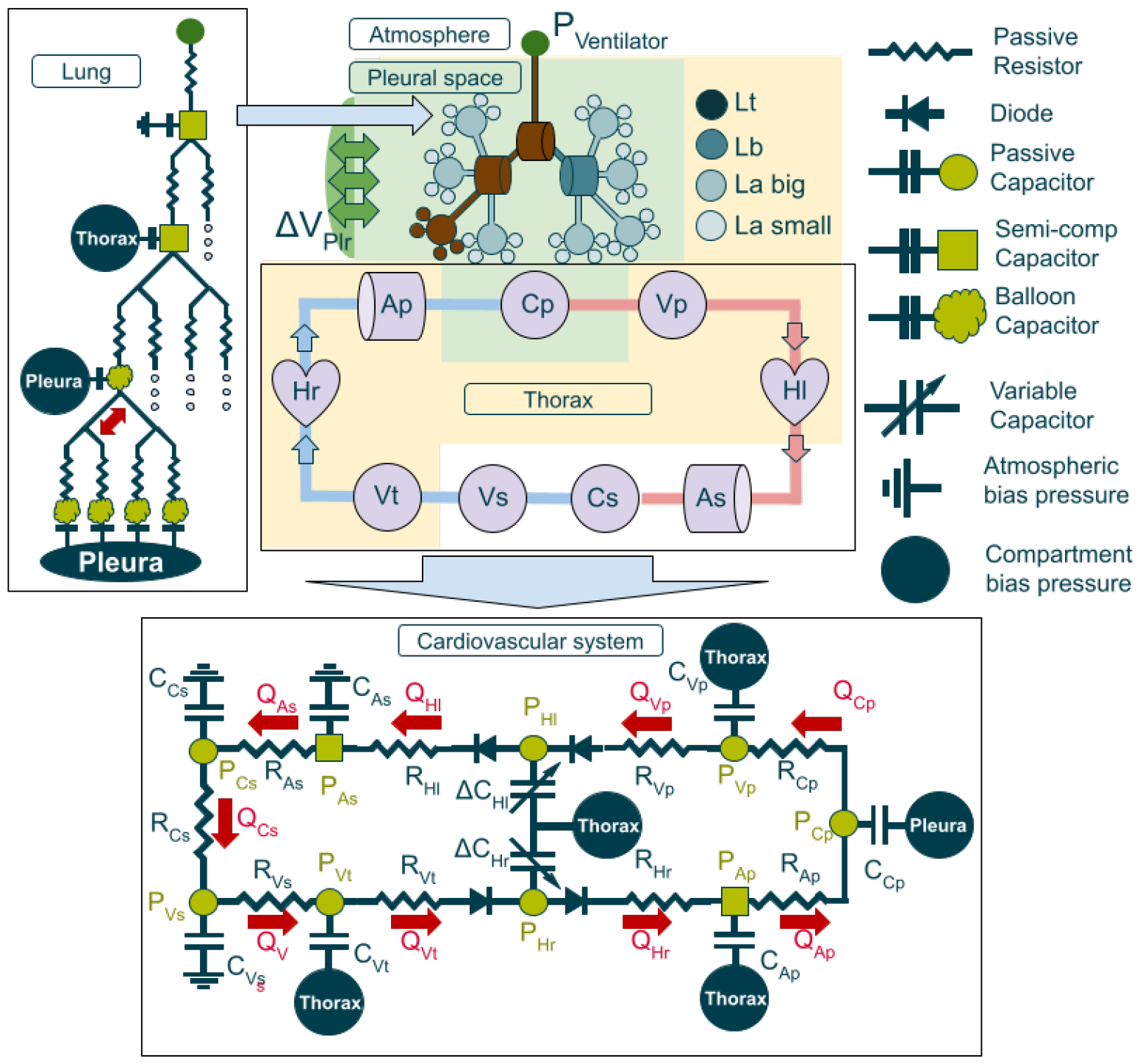
Schematic representation of the CP model with the equivalent electric circuits for the pulmonary and cardiovascular systems.

#### Vasculature

The vasculature refers to the network of blood vessels that circulate blood throughout the body, including arteries, veins, and capillaries. Since arteries are less compliant than veins, these vessels would therefore need more pressure to be exerted to push the same amount of fluid into the compartment. This is particularly relevant for vessels with thicker walls, where the force exerted on the vessel wall relative to its deformation from its original shape generates a significant counter-reaction. The constant *V*_0_ in (1) is used to account for this effect, enabling the model to accurately represent distinct compliance characteristics of different types of blood vessels. The parameter *V*_0_ can be seen as the unstressed volume, i.e. the equilibrium volume the vessel would have when the pressure in the system is equal to the bias pressure [18]. The appropriate choice of *V*_0_ allows to impart the arterial system with desired blood volume whilst being able to maintain low compliance on the system so that high pressures can be attained without significant changes to the volume.

#### Valves

The valves in the circulatory system essentially act as gates, ensuring the one-way flow of blood to maintain proper circulation.

Valve dynamics are modeled using

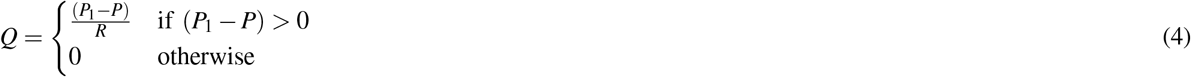

where *P*_1_ and *P* are the pressures across the valve. The valves block the flow and resets *Q* to zero whenever the pressure difference becomes negative in the compartment.

#### Heart

The heart functions as a muscle that operates in a periodic cycle of contraction (systole) and relaxation (diastole). Consequently, the compliances representing the heart chambers also have to be specifically tailored to account for the cyclic nature of cardiac function. In literature, the most common approach is to modulate the value of compliance using elastance models. We use the variable-elastance model adapted from Heldt et al. [34] written as

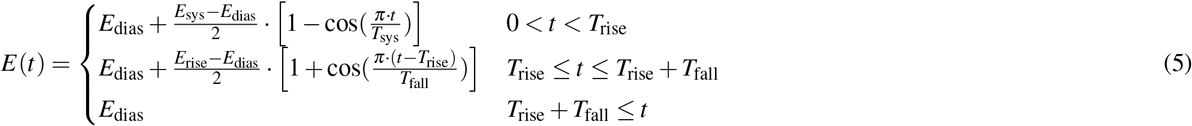

The compliance is then computed from equation (5) as *C*(*t*) = 1*/E*(*t*) and is subsequently used in equation (1). Here *t* denotes time measured with respect to the onset of ventricular contraction. *E*_sys_ and *E*_dias_ represent the values for the systolic and diastolic elastances respectively. *T*_rise_ and *T*_fall_ is the duration of the rise and fall of the elastance profile during the systole and diastole respectively. The parameter *E*_sys_, is instrumental in modulating the intensity of the heart’s contractions. Similarly, *E*_dias_ modulates the volume of blood that is allowed back into the heart during the diastolic phase. Thus the elastance equation (5) governs the ventricular activation, which is pivotal in maintaining blood flow across all compartments throughout a cardiac cycle.

#### Lungs

The respiratory model structure consists of a trachea bifurcating into two bronchi, which further branch out into a tree-like network of alveoli, simulating the intricate architecture of the human respiratory system. In contrast to the CV system, the PS is an open-loop system that is responsible for transporting air from the atmosphere through the trachea and upper airways into the alveoli where oxygen and carbon dioxide are exchanged with blood in the pulmonary capillaries. The PS is connected to the atmosphere through the pharynx, extends through the throat and trachea, and is subsequently distributed through the lung via a network of bronchi terminating in the alveolar region. The mathematical formulation follows the same principles as the ones described for the CV components. The trachea and the bronchi are airways cladded with cartilage and smooth muscle and therefore their compliance is expected to be very low. These airways, except for the bronchioles, are not supposed to collapse and will hold most of their volume as unstressed volumes for these reasons; equation (1) is used to model the pressure changes in these compartments. The alveoli experiences significant volume change during normal respiration which leads to a non-linear relationship between its compliance and volume. A key aspect of this relationship is the concept of lung collapse, which primarily occurs at the level of individual alveoli. This phenomenon takes place when the air volume within an alveolus drops sufficiently low, and the transpulmonary pressure between the pleura and the alveoli is insufficient to counteract the elastic and the surface tension forces on the alveolar walls. Under these conditions, the alveoli lose their connection to the bronchi and become incapable of engaging in gas exchange. Reopening a collapsed alveolus requires exerting a specific amount of pressure referred to as the Threshold Opening Pressure (TOP) to overcome the surface tension, allowing the alveolus to ‘pop’ open again. This reopening process is essential for restoring normal alveolar function and ensuring effective gas exchange [14].

The aforementioned alveoli dynamics share a lot of the same characteristics with rubber balloons, where creating the initial extension requires the transmural pressure to exceed a certain threshold value, beyond this point, the pressure required to keep inflating the balloon decreases until it starts overstretching resulting in increasingly higher pressures being necessary to inflate any further. The mathematical formulation of spherical elastic balloon dynamics can be found in the works of Muller et al [35]. Of particular interest to this work is the pressure/radius curve presented which is analogous to a pressure/volume curve in our model. A similar pressure/volume behaviour can be obtained when the compliance of the compartment is calculated by merging a reversed sigmoid curve (equation 6) with another sigmoid curve (equation 7) as shown in Figure 3.

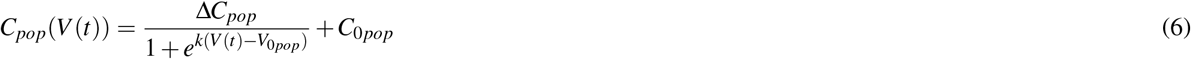

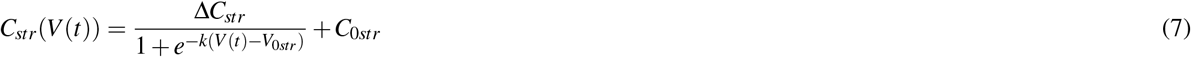

The resulting pressure can be obtained by calculating *P*(*t*) = *V* (*t*)*/C*(*V* (*t*)). This pressure refers to the transmural pressure that must exceed a certain threshold value to maintain the alveoli in an open state. The compliance curve produced as a result creates a pressure profile that imparts a non-linear behavior to the system, characterized by four distinct operational modes. The first mode is a closed mode that operates between 0mL and the volume at the peak of the pressure curve (TOP), where very small variations of volume generate high changes in pressure. The second mode of operation lies between the volume at TOP and the volume at the local pressure minimum, here the alveoli are considered to be opening in a cascade effect generated by the drop in transmural pressure needed to keep the alveoli open. The third mode of operation is characterised by a relatively small increase in pressure needed to keep inflating the alveoli. The fourth mode occurs beyond a point in volume where the inflection point of the stretching sigmoid is located, in this regime the pressure/volume relationship starts to be exponential and the alveoli are considered to have reached maximum expansion and begin to undergo tissue over-stretching. Consequently, the volume at this stage can be regarded as being at, or close to, the maximum capacity that the compartment can sustain under physiological pressures.

**Figure 3.**
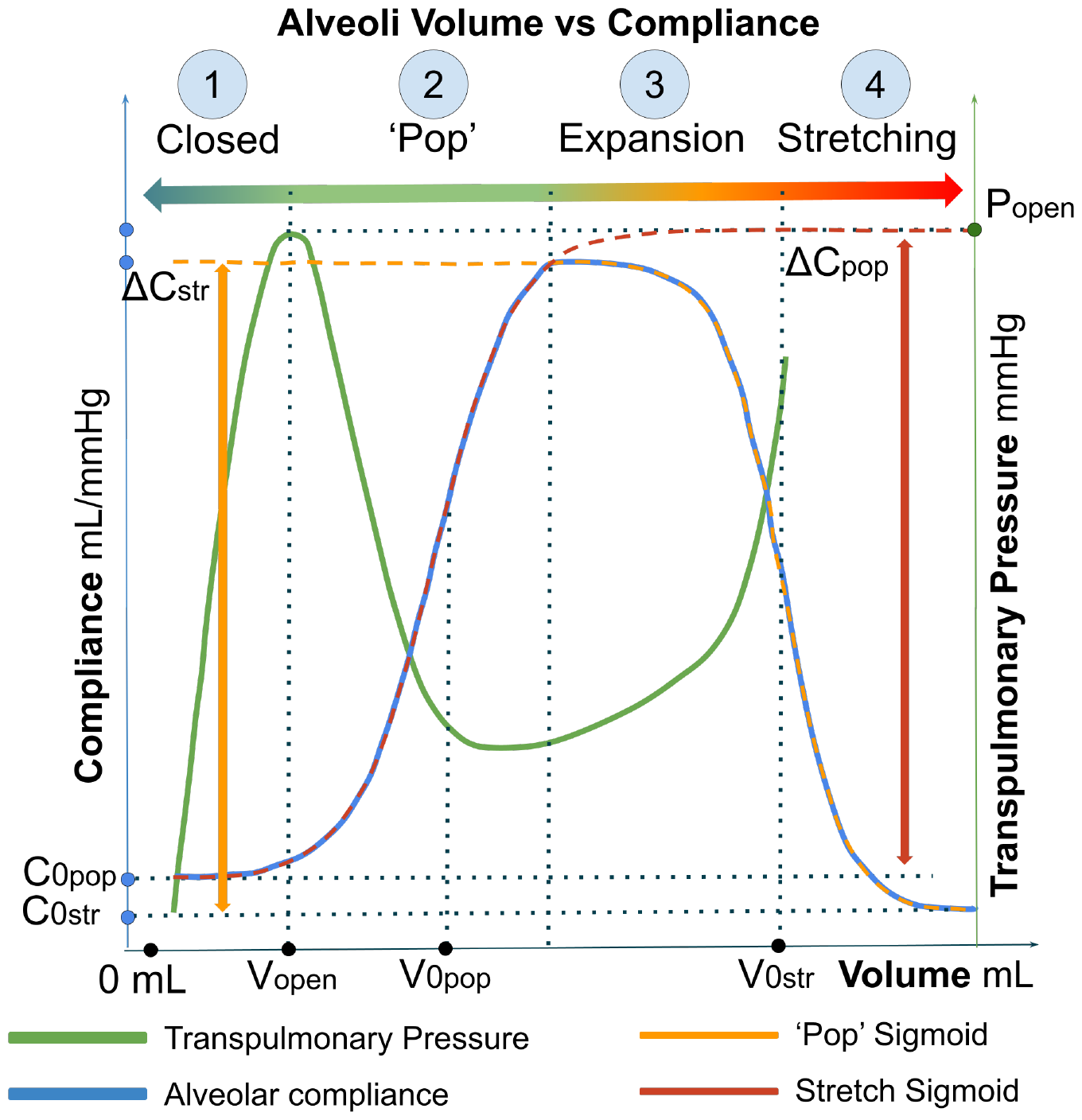
Compliance-volume relation of a balloon alveoli with four distinct operational modes of an alveoli highlighted. **Mode 1:** Closed Mode, ranging from 0*mL* to the volume at the peak of the pressure curve (TOP), where small volume changes result in significant pressure variations. **Mode 2**: Cascade opening Mode (‘Pop’), occurring between the volume at TOP and the volume at the local pressure minimum, characterized by alveoli opening due to reduced transmural pressure. **Mode 3:** Expansion Mode, where a small increase in pressure leads to further alveolar inflation, indicating the lung’s expansion phase. **Mode 4:** Over-Stretching Mode, beyond the inflection point of the stretching sigmoid, marking the onset of exponential pressure/volume relationship and maximal alveolar expansion, leading to potential tissue over-stretching. This final mode represents the threshold of maximum compartment volume achievable at physiological pressures.

This novel strategy introduces a more realistic method for modelling alveoli, incorporating non-linear alveolar dynamics and lung heterogeneity. Contrary to traditional models in the literature that depict alveolar opening as a step increase in volume after reaching a critical TOP, our model simulates alveolar opening as a process that emerges from a time-dependent compliance/volume relation. This approach not only offers a more precise representation of alveolar states but also delineates four distinct operating zones for the alveoli. The proposed equations to model alveolar dynamics include eight distinct parameters for each alveolus. The parameter *C*_0*pop*_, in equation (6), specifies the alveolus’s residual volume when it is in a closed state. Similarly, *C*_0*str*_ from equation (7) determines the alveoli’s residual compliance when it is overstretched, influencing the steepness of the pressure curve in this condition. The combination of Δ*C*_*pop*_, the slope parameter *k*, and *V*_0*pop*_ establishes the TOP, with *V*_0*pop*_ being crucial in determining the opening volume of the alveoli (typically a few mL below *V*_0*pop*_). *V*_0*str*_ defines the volume at the inflection point on the stretching sigmoidal curve, limiting the maximum volume of air allowed in the alveoli.

The branched architecture of the PS is created using a recursive tree generation algorithm, enabling an efficient construction of a lung model with desired number of alveolar units. This method ensures the model’s adaptability to various scenarios while maintaining computational efficiency for real-time simulations. The PS model used to produced in the results presented consists of a single trachea compartment, two bronchi compartments, and forty alveoli, with each bronchus dividing into four alveoli, and each alveolus further branching into four additional alveoli. This structure results in eight groups of five alveoli, where four of them are directly connected to a parent alveolus. To mimic realistic physiological variations, each alveolus is assigned a slightly different TOP, allowing for heterogeneity during simulations. Figure 4 illustrates the compliance/volume and pressure/volume relationships for some selected alveoli, offering insight into the dynamics of alveolar opening and its effects on pressure and volume variations across the PS.

**Figure 4.**
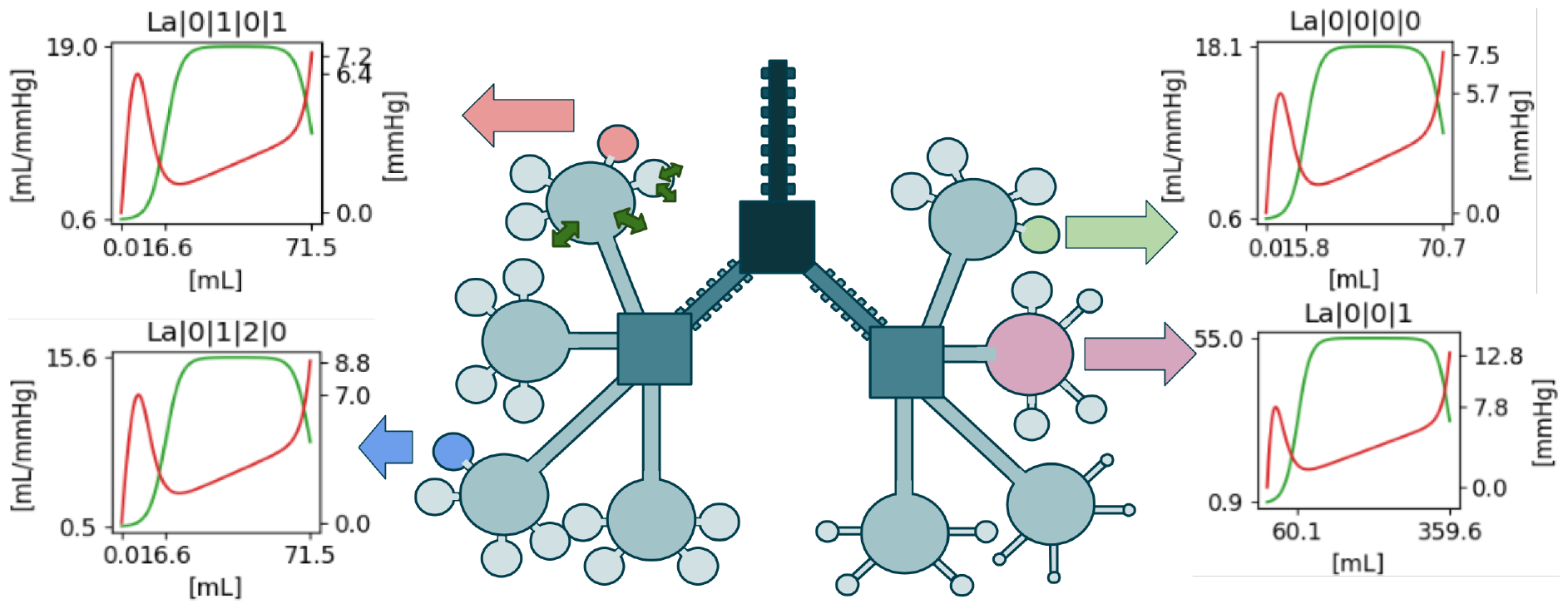
Visualization of the compliance/volume profiles (shown in green) and pressure/volume profiles (shown in red) for selected alveoli, highlighting their nonlinear behavior and the resulting heterogeneity of the lung.

#### Thoracic cavity

One of the challenges in modeling the CP system using zero-dimensional models is to address the influence of surrounding regions on each compartment. Two critical regions in this context are the thoracic cavity and the pleural space. The thoracic cavity is the region enclosed by the rib cage and the diaphragm encompassing the heart along with its inflow and outflow vessels, as well as the respiratory system. The pleural cavity is the liquid-filled region between the parietal pleura (lining the thoracic cavity) and the visceral pleura (covering the lungs) that envelopes the lungs and all the blood vessels that irrigate it. The parameter *P*_*bias*_ in equation (1) is employed to account for the bias pressure that different compartments experience, based on their spatial location. In our model, we use the pressures at the pleura and thoracic cavity to establish the bias pressures at the specific locations indicated in Figure 2. The thoracic cavity is modeled as a semi-compliant capacitor where *V* from equation (1) is obtained by aggregating the volumes of all compartments encompassed within the thoracic cavity. *V*_0_ denotes the unstressed volume of the thoracic cavity, functioning as the diaphragm’s activation mechanism, with the thoracic cavity’s pressure being biased by atmospheric conditions. Since the *P*_bias_ is consistent for compartments situated within the same cavity, any changes in pressure within one compartment are effectively transmitted to the other compartments via the thoracic pressure.

#### Spontaneous Breathing and Mechanical Ventilation

In our model, the spontaneous breathing mechanism is simulated using a methodology akin to that used for modeling cardiac muscle activity. We achieve this by adapting the temporal dynamics of the unstressed thoracic volume throughout a respiratory cycle RC, applying specific modifications to (5). Specifically, the *E*_sys_ and *E*_dias_ in the cardiac activation function are replaced with *V* ^max^ and *V* ^min^ respectively. Furthermore, *T*_rise_ is changed to *T*_insp_ and set to 0.33 *· RC*, where *RC* is calculated from the respiratory rate as *RR* = 1*/*(60 *×RC*). The modified equation is then written as

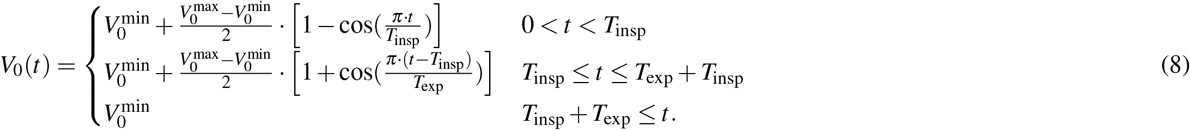

The unstressed volume *V*_0_ is then used to calculate the thoracic pressure using (1). In the case of mechanical ventilation, it is common for patients to be either not spontaneously breathing or under an assisted breathing regimen provided by a ventilator. Our model simulates these conditions by modulating air pressure at the mouth as follows:

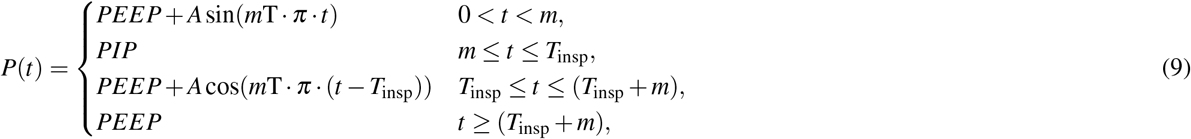

where *t* is the relative time computed from the beginning of an RC, *PEEP* is the peak end-expiratory pressure, *PIP* is the peak inspiratory pressure, *T*_insp_ is the duration of inspiration^1^, *m* is the transition time between expiration/inspiration and *A* is the pressure amplitude. This equation generates a square wave with rounded edges and simulates a mechanical ventilator operating in pressure-control mode, where a minimum PEEP above atmospheric pressure is constantly maintained. To enable an inspiration, the ventilator increases the pressure to PIP.

### Computer implementation of the CP model

The CP model architecture, depicted in Figure 2, is configured by assembling a number of generic compartments. This construction is facilitated by encoding the model’s graph structure into a JavaScript Object Notation (JSON) format, derived from the data across Tables 1 to 5, to parameterize and initialise the system of governing ODEs of our model. The implementation was executed in Python 3.1, augmented with the Just After eXecution (JAX) library for its efficiency in numerical computations, native graphics processing unit (GPU) acceleration, and Just in Time (JIT) compilation capabilities, enhancing computational performance. The Equinox library was used for defining and managing equation classes and the integration stack, owing to its seamless integration with JAX and transformation of parameterized functions into PyTrees. Additionally, the Diffrax library, another JAX-based tool, was employed for its advanced numerical differential equation solving capabilities. Numerical integration was performed using Euler’s method with a fixed step size of 0.01 seconds on a high-specification CPU (Intel Core i7-8565U with 32GB RAM), without GPU utilization. The algorithm begins by computing the pressures in the regions (Table 4) followed by pressures in compartments (Table 2), and flow rates (*Q*) across resistors (Table 1). Subsequently, the temporal evolution of volume *dV/dt* for all compartments is evaluated using the equations of Table 3. The output from this model comprises high-fidelity (100Hz) time-series data of pressures, volumes, and flow within each compartment and flow rates between different compartments. This efficient computer implementation, equipped with Python-based libraries such as JAX and Diffrax, ensures the fast generation of high-frequency data in real time, essential for the in-depth CP system analysis required from a digital twin environment.

**Table 1.**
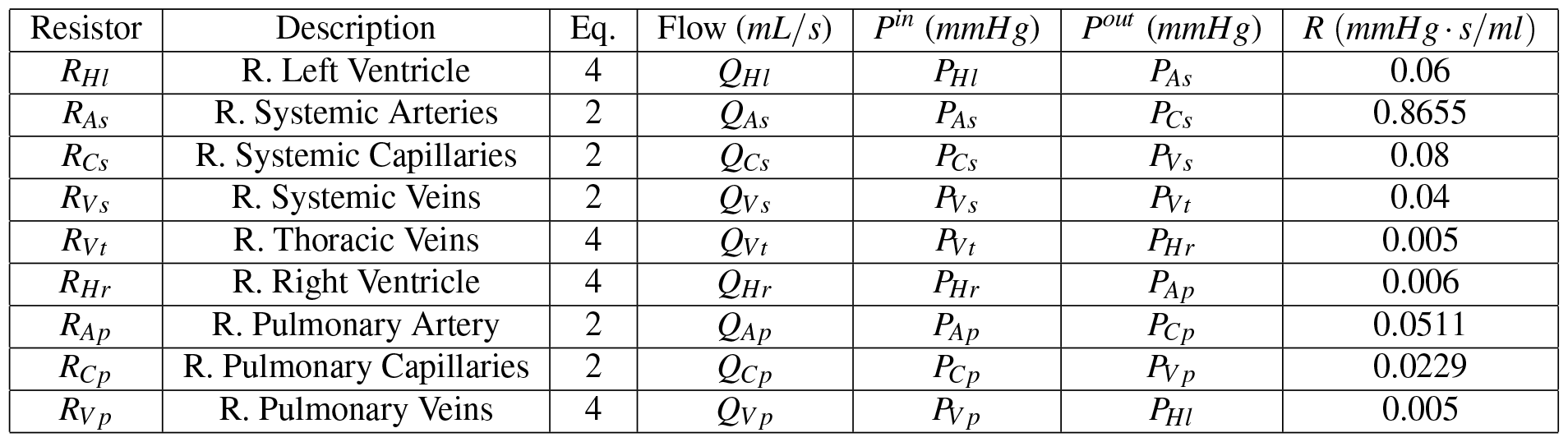
Generic data structure used to calculate the flows across different compartments in the CP model. The flow between any two compartments denoted in column 4 is determined by dividing the pressure difference across these compartments listed in the columns *P*^*in*^ and *P*^*out*^ by the specific resistance type that exists between them. The column Eq.shows the model equation that calculates the flow based on the type of resistor.

| Resistor | Description | Eq. | Flow (mL/s) | $P^{in}$ (mmHg) | $P^{out}$ (mmHg) | $R$ (mmHg · s/ml) |
| --- | --- | --- | --- | --- | --- | --- |
| $R_{Hl}$ | R. Left Ventricle | 4 | $Q_{Hl}$ | $P_{Hl}$ | $P_{As}$ | 0.06 |
| $R_{As}$ | R. Systemic Arteries | 2 | $Q_{As}$ | $P_{As}$ | $P_{Cs}$ | 0.8655 |
| $R_{Cs}$ | R. Systemic Capillaries | 2 | $Q_{Cs}$ | $P_{Cs}$ | $P_{Vs}$ | 0.08 |
| $R_{Vs}$ | R. Systemic Veins | 2 | $Q_{Vs}$ | $P_{Vs}$ | $P_{Vt}$ | 0.04 |
| $R_{Vt}$ | R. Thoracic Veins | 4 | $Q_{Vt}$ | $P_{Vt}$ | $P_{Hr}$ | 0.005 |
| $R_{Hr}$ | R. Right Ventricle | 4 | $Q_{Hr}$ | $P_{Hr}$ | $P_{Ap}$ | 0.006 |
| $R_{Ap}$ | R. Pulmonary Artery | 2 | $Q_{Ap}$ | $P_{Ap}$ | $P_{Cp}$ | 0.0511 |
| $R_{Cp}$ | R. Pulmonary Capillaries | 2 | $Q_{Cp}$ | $P_{Cp}$ | $P_{Vp}$ | 0.0229 |
| $R_{Vp}$ | R. Pulmonary Veins | 4 | $Q_{Vp}$ | $P_{Vp}$ | $P_{Hl}$ | 0.005 |

**Table 2.**
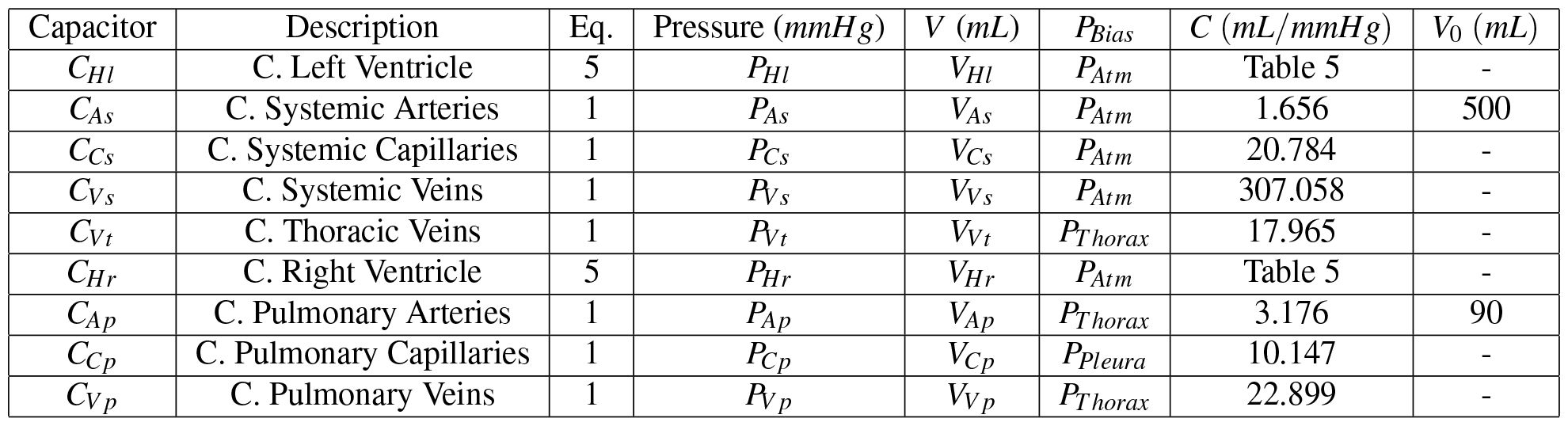
Generic data structure used for the calculation of pressures in a compartment based on the different types of capacitors utilized in the model. The pressure in any compartment is determined by dividing the instantaneous volume in that compartment given by column “*V*” and the unstressed volume (*V*_0_) by the specific type of capacitor of that compartment.

| Capacitor | Description | Eq. | Pressure (mmHg) | $V$ (mL) | $P_{Bias}$ | $C$ (mL/mmHg) | $V_0$ (mL) |
| --- | --- | --- | --- | --- | --- | --- | --- |
| $C_{Hl}$ | C. Left Ventricle | 5 | $P_{Hl}$ | $V_{Hl}$ | $P_{Atm}$ | Table 5 | - |
| $C_{As}$ | C. Systemic Arteries | 1 | $P_{As}$ | $V_{As}$ | $P_{Atm}$ | 1.656 | 500 |
| $C_{Cs}$ | C. Systemic Capillaries | 1 | $P_{Cs}$ | $V_{Cs}$ | $P_{Atm}$ | 20.784 | - |
| $C_{Vs}$ | C. Systemic Veins | 1 | $P_{Vs}$ | $V_{Vs}$ | $P_{Atm}$ | 307.058 | - |
| $C_{Vt}$ | C. Thoracic Veins | 1 | $P_{Vt}$ | $V_{Vt}$ | $P_{Thorax}$ | 17.965 | - |
| $C_{Hr}$ | C. Right Ventricle | 5 | $P_{Hr}$ | $V_{Hr}$ | $P_{Atm}$ | Table 5 | - |
| $C_{Ap}$ | C. Pulmonary Arteries | 1 | $P_{Ap}$ | $V_{Ap}$ | $P_{Thorax}$ | 3.176 | 90 |
| $C_{Cp}$ | C. Pulmonary Capillaries | 1 | $P_{Cp}$ | $V_{Cp}$ | $P_{Pleura}$ | 10.147 | - |
| $C_{Vp}$ | C. Pulmonary Veins | 1 | $P_{Vp}$ | $V_{Vp}$ | $P_{Thorax}$ | 22.899 | - |

**Table 3.** Calculation of the temporal evolution of volume in each compartment in the model. The volume change is determined by the resistances that modulate the flow in and out of the compartment listed in columns *R*^*in*^ and *R*^*out*^ and the compliance of that compartment listed in the Capacitor column. The model equation that computes the volume is specified in the last column.

| Comp | Description | Capacitor | $R^{in}$ | $R^{out}$ | $dV/dt$ | eq. |
| --- | --- | --- | --- | --- | --- | --- |
| $Hl$ | Left Ventricle | $C_{Hl}$ | $R_{Hl}$ | $R_{As}$ | $dV_{Hl}/dt$ | 3 |
| $As$ | Systemic Arteries | $C_{As}$ | $R_{As}$ | $R_{Cs}$ | $dV_{As}/dt$ | 3 |
| $Cs$ | Systemic Capillaries | $C_{Cs}$ | $R_{Cs}$ | $R_{Vs}$ | $dV_{Cs}/dt$ | 3 |
| $Vs$ | Systemic Veins | $C_{Vs}$ | $R_{Vs}$ | $R_{Vt}$ | $dV_{Vs}/dt$ | 3 |
| $Vt$ | Thoracic Veins | $C_{Vt}$ | $R_{Vt}$ | $R_{Hr}$ | $dV_{Vt}/dt$ | 3 |
| $Hr$ | Right Ventricle | $C_{Hr}$ | $R_{Hr}$ | $R_{Ap}$ | $dV_{Hr}/dt$ | 3 |
| $Ap$ | Pulmonary Arteries | $C_{Ap}$ | $R_{Ap}$ | $R_{Cp}$ | $dV_{Ap}/dt$ | 3 |
| $Cp$ | Pulmonary Capillaries | $C_{Cp}$ | $R_{Cp}$ | $R_{Vp}$ | $dV_{Cp}/dt$ | 3 |
| $Vp$ | Pulmonary Veins | $C_{Vp}$ | $R_{Vp}$ | $R_{Hl}$ | $dV_{Vp}/dt$ | 3 |

**Table 4.** Calculation of the total volumes of the thorax and pleura regions. The total volume in these compartments is essentially the sum of the volumes of all the compartments inside these regions.

| Name | Description | Eq. | Volumes (mL) | $P_{bias}$ (mmHg) | $C$ (mL/mmHg) | $V_0$ (mL) |
| --- | --- | --- | --- | --- | --- | --- |
| *Thorax* | Thoracic pressure | 1 | $V_{Ap} + V_{Cp} + V_{Vp} + V_{Hl} + V_{Hr} + V_{Vt}$ | $P_{Atm}$ | 300 | 3500 |
| *Pleura* | Pleural pressure | 1,8 | $V_{Ap} + V_{Cp} + V_{Vp}$ | $P_{Atm}$ | 35 | Table 5 |

### Model Parameters

Although our CP model is influenced by the modelling approaches found in existing literature [17, 25], substantial modifications to the model architecture and the incorporation of novel alveolar dynamics, necessitate a modification to the parameters from these studies. Consequently, to accurately calibrate our model, we implement an optimisation-based approach for parameter evaluation. Our calibration procedure aims to ensure that the model accurately captures physiological responses while operating within a normal range. Specifically, we strive to simulate the physiological behaviour of a healthy human, characterized by typical volume distribution and pressure levels throughout the CP system, as detailed in Tables 7 and 8. In essence, the calibration process involves adjusting the compliance and resistance values within the system to achieve these set target ranges of pressure and volume.

#### CV optimisation strategy

On the cardiovascular side, the parameters proposed for resistance and compliance/elastance in [34] were used as the initial values in the calibration process. From these, eleven were selected for optimisation, leaving the remainder unchanged. The parameters chosen were the two resistors for the arterial systems, *R*_*As*_ and *R*_*Ap*_, the systolic elastances *E*_*sys*_*Hl* and *E*_*sys*_*Hr* and all seven compliances *C* of all compartments representing blood vessels.

From (1) it can be observed that the value for compliance in a compartment affects the pressure and volume of a compartment. If the compliance of a compartment goes up, the pressure in that compartment will go down (negatively correlated) and the volume held in the compartment will go up (positively correlated). Using the same idea, it can be seen from (2) that the value for the resistance connecting two compartments modulates the flow rate that is allowed between these compartments, where higher resistance for the same pressure difference will yield less flow rate.

Knowledge of these relations between the parameters and the variables can be leveraged to create an extra set of differential equations that will nudge the model to settle into physiologically relevant zones where the values for pressure and volume generated by the model can be targeted. To illustrate our parameterization methodology, take the systemic capillaries as an example, where in Guyton et al. [36] it is stated that the normal volume of blood in these vessels is 7% of the total blood volume. To nudge the model into settling in this volume another equation 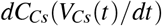 is added to the model. The equations operate by, firstly, calculating the relative error between the volumes observed and the target volume *σ* = (*V*_current_*/V*_target_) *−* 1. If the volume is below the target, compliance needs to increase and the magnitude of the increase/decrease of the value of the compliance should be proportional to the residual error. If the variables are anti-correlated, the relationship inverts. The variation of compliance is calculated as

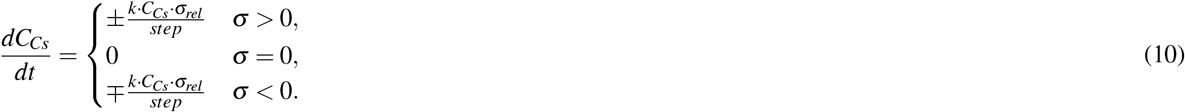

where *k* is a constant that modulates the magnitude of change for compliance.

A *C*_*min*_ and a *C*_*max*_ are used to prevent the system constrained to reasonable values. Table 6 are all the equations added to the model during the calibration process with the respective targets. The equation used to calibrate the value for *C*_*Vs*_ was made to target a pressure instead of a volume because the same volume distribution in the system can be achieved for different operating pressures: a pressure target, therefore, better directed the system to settle within a specified pressure range. The stroke volume used as a target for the Resistor was calculated by integrating the flow between the heart and the artery and updated at the end of every heartbeat and the systemic and pulmonary arterial pressures were used to modulate the systolic elastance, with the diastolic elastance left constant according to [34].

The system was then allowed to run until it reached equilibrium and the resulting parameters are presented in Tables 1, 2 and 5

**Table 5.** Base data structure that parameterises a variable capacitor and the ventilator function.

| Name | Description | $E_{max}$ (mmHg/mL) | $E_{min}$ (mmHg/mL) | $T_{sys}$ (s) | Cycle Period (s) | eq. |
| --- | --- | --- | --- | --- | --- | --- |
| $E_{Hl}$ | Left Ventricle Elastance | 2.812 | 0.0686 | $0.3\sqrt{0.8}$ | 0.8 | 5 |
| $E_{Hr}$ | Right Ventricle Elastance | 0.575 | 0.0414 | $0.3\sqrt{0.8}$ | 0.8 | 5 |
| $V_{0Pleura}$ | Pleura Unstressed Volume | 3850 | 3300 | 0.2 | 5 | 8 |
| Name | Description | $PEEP$ (mmHg) | $amplitude$ (mmHg) | $T_{insp}$ (s) | Cycle Period (s) | eq. |
| $P_{Vent}$ | Ventilator Pressure | 5 | 8 | 0.2 | 5 | 9 |

**Table 6.** Descriptors for the equations used in the calibration process.

| Target Variable | Target | Parameter to calibrate | +/- | Min | Max |
| --- | --- | --- | --- | --- | --- |
| $V_{As}$ | 650 mL | $C_{As}$ | + | 0.01 | 10 |
| $V_{Cs}$ | 350 mL | $C_{Cs}$ | + | 0.01 | 50 |
| $P_{Vs}$ | 10 mmHg | $C_{Vs}$ | - | 150 | 1400 |
| $V_{Vt}$ | 100 mL | $C_{Vt}$ | + | 0.01 | 50 |
| $V_{Ap}$ | 150 mL | $C_{Ap}$ | + | 0.01 | 10 |
| $V_{Cp}$ | 150 mL | $C_{Cp}$ | + | 0.01 | 15 |
| $V_{Vp}$ | 300 mL | $C_{Vp}$ | + | 0.01 | 50 |
| $SV_{Hr}$ | 68 mL | $R_{Ap}$ | - | 0.01 | 200 |
| $SV_{Hl}$ | 68 mL | $R_{As}$ | - | 0.01 | 200 |
| $P_{As}$ | 90 mmHg | $E_{maxHl}$ | + | 0.01 | 4 |
| $P_{Ap}$ | 17 mmHg | $E_{maxHr}$ | + | 0.01 | 1 |

**Table 7.** Simulated pressures at every compartment compared to the normal values observed in these compartments found in literature [34] over the 10-second simulation period.

| Compartment | Pressure (mmHg) |  |  |  |  |
| --- | --- | --- | --- | --- | --- |
|  | Min | Avg | Max | Diast/Min [34] | Syst/Max [34] |
| Systemic Artery | 75.5 | 95.7 | 115.2 | 60-90 | 90-140 |
| Systemic Capillaries | 16.7 | 16.9 | 17.1 | - | - |
| Systemic Veins | 9.6 | 9.6 | 9.7 | - | - |
| Thoracic Veins | 5.1 | 6.0 | 7.3 | 0-8 | 2-14 |
| Right Ventricle | 2.5 | 11.5 | 31.3 | 0-8 | 15-28 |
| Pulmonary Artery | 13.8 | 19.9 | 29.6 | 5-16 | 15-28 |
| Pulmonary Capillaries | 13.0 | 17.0 | 23.5 | 1-12 | 9-23 |
| Pulmonary Veins | 11.7 | 13.1 | 15.7 | - | - |
| Left Ventricle | 4.9 | 40.0 | 116.0 | 4-12 | 90-140 |

**Table 8.**
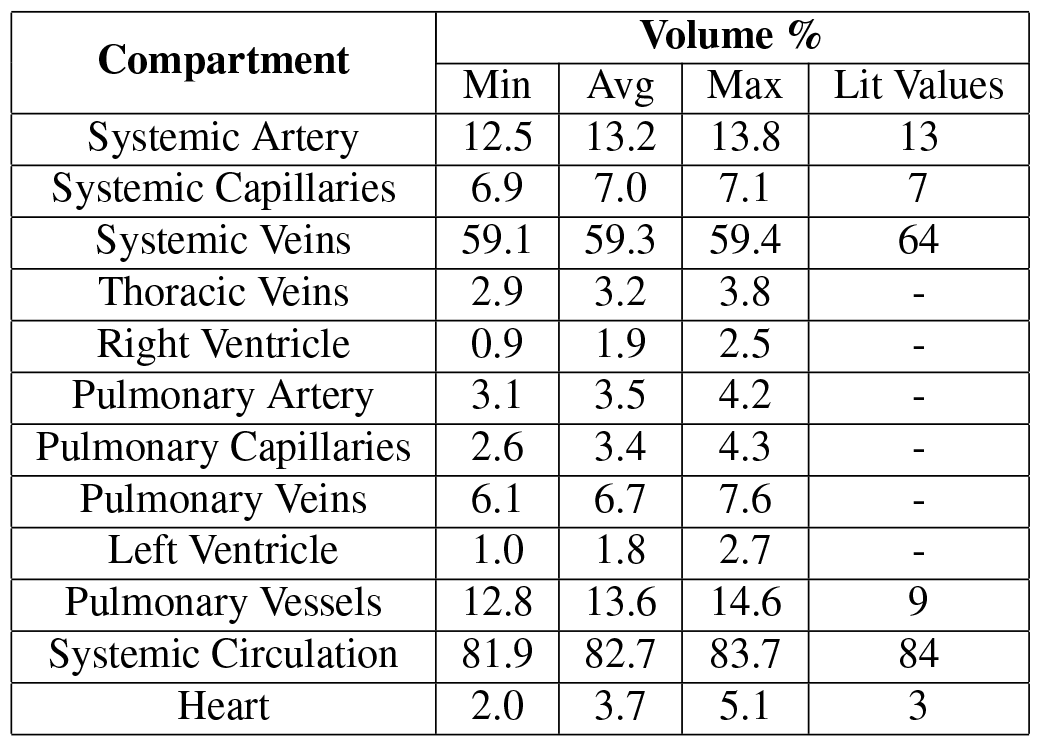
Simulated Volumes at every compartment compared to the normal values observed in these compartments found in literature [36] over the 10-second simulation period.

#### PS optimisation strategy

The complexity of the chosen PS model prevented us from using a similar technique to parametrise the model. Moreover, due to the novelty of the strategy used, there were no literature values that could be leveraged for alveoli parametrisation and a first principle-based approach had to be employed instead, targeting normal lung volumes. Please see in [36] (Table 10).

The trachea and both bronchi are the compartments responsible for most of the respiratory dead space and serve mainly as resistive vessels with small volume variations. To model this they were initially given very low compliance (0.5 and 2 mL/mmHg) and resistance (0.002 and 0.003 *mmHg· s/mL*) respectively. To modulate the baseline volume in these compartments a total unstressed volume of 225 mL (3 *×* 75 mL) was attributed.

The interpretability of the parameters of the alveoli equations (6-7) was of paramount importance in the parameterization process of the pulmonary system. The value for the sum of all *V*_0*str*_ across all alveoli is always very close to the maximum volume of the pulmonary system. *V*_0*pop*_ and Δ*C*_*pop*_ are the main determinants of TOP and tiny variations of these parameters will generate different TOP in every alveoli with *k* determining the steepness of the pressure wall. This flexibility allows the pulmonary system to be re-parametrized to represent different lung dynamics with a wide range of opening profiles.

The 8 parent alveoli were given a higher inflow resistance of 0.03 *mmHg · s/mL* to reflect the fact that the highest lung resistance resides in the bronchioles and the 32 child alveoli were given a lower resistance of 0.003 *mmHg ·s/mL*. To model a Functional Residual Capacity FRC of 2500 *mL* the total *V*_0*str*_ of the 8 parent alveoli was set to 312.5 *mL* and to ensure that these alveoli open at very low volumes *k* = 0.1m^*−*3^, *V*_0*pop*_ = 60*mL*, Δ*C*_*str/pop*_ = 55*mL/mmHg* and *C*_0*str/pop*_ = 0.5*mL/mmHg* generating a TOP of 7.8 *mmHg*.

The parameters used for the child alveoli were *k* = 0.35 and *C*_0*str/pop*_ = 0.5*mL/mmHg*, and to create a heterogeneous lung *V*_0*pop*_ and Δ*C*_*str/pop*_ were set to 17*mL ±* 1.5 and 17.0*mL/mmHg ±* 3 using a normally distributed random number generator. To create a total lung capacity of around 5000 *mL, V*_0*str*_ was set to 55 *mL* + *V*_0*pop*_. This created a set of alveoli with TOPs ranging from 5.4 *mmHg* and 8.2 *mmHg*. In Figure 4, the properties of some alveoli examples are presented.

## 3 RESULTS

### Baseline simulation

To assess the suitability of the model to represent the CV dynamics and CP interactions of a healthy human, a baseline simulation was performed where the system was allowed to stabilise while spontaneously breathing at 12 *breaths/minute* with a heart rate of 75 *beats/minute*. Figure 5 presents the distribution of blood volume and pressures across the CV compartments. The results are presented over two RC (10 seconds). In this scenario, the system displays a mean systemic arterial blood pressure BP of 91.16 mmHg (75.52-115.25 diastolic and systolic) and an average stroke volume SV of 72.7mL, generating a CO of 5.452 L/min.

**Figure 5.**
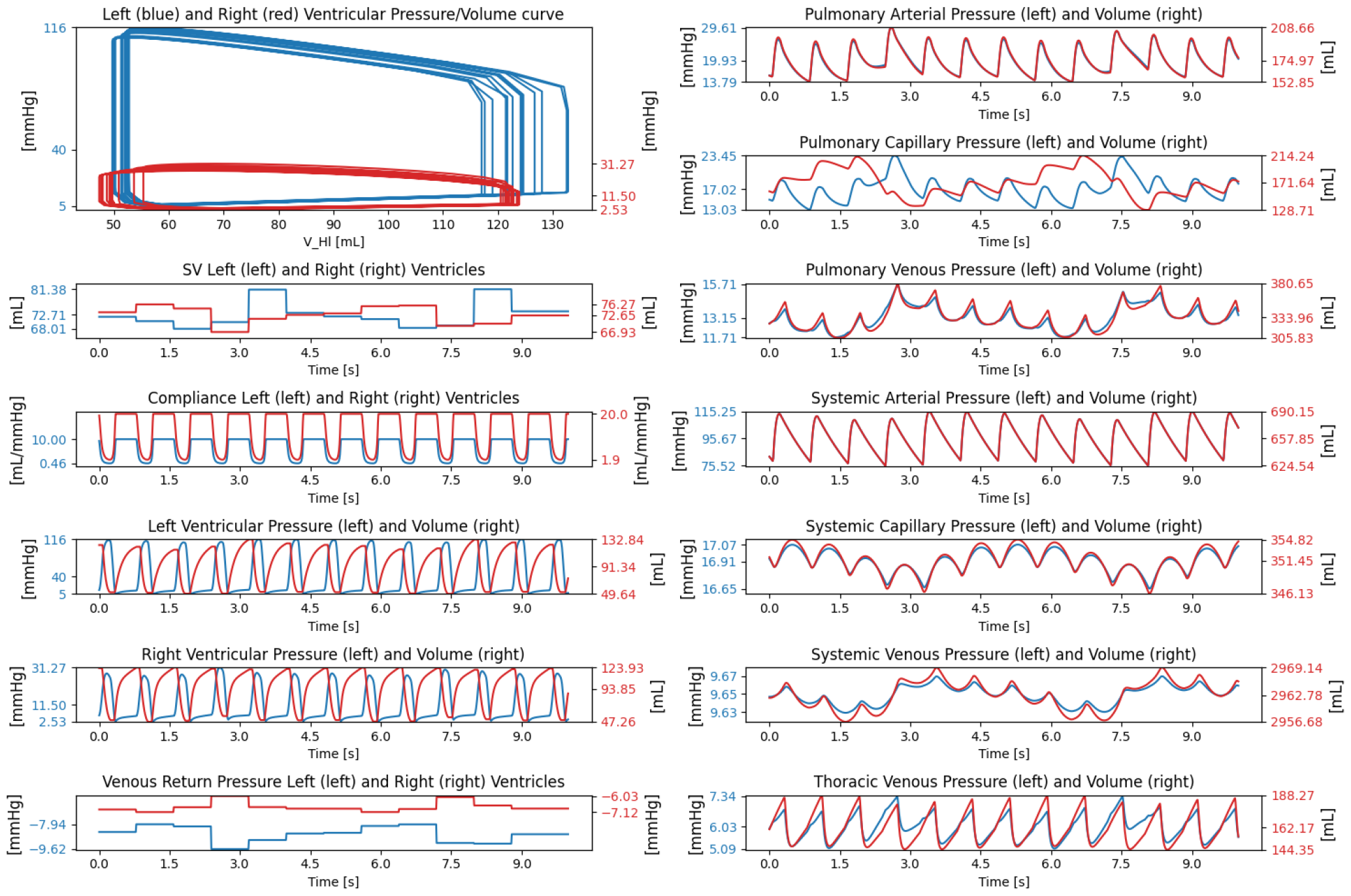
Baseline Simulation of the cardiovascular portion of the model over two full spontaneous breaths. Maximum, average and minimum values observed during simulation are labelled on vertical axes.

The simplicity of the model equations, the parameter interpretability and the graph nature of the system provided useful guidance during the parameterization process, allowing us to identify the important parameters that ensure the model is operating in a regime considered representative of normal human physiology. Tables 7 and 8 present a comparison between the simulated pressures and volumes with the values usually observed in healthy humans. As shown, the mean values obtained for the volumes and pressures of the cardiovascular compartments where the ‘optimisers’ are used, are all within the targets defined during the optimization process.

#### Self-breathing

The lower frequency oscillation of the BPs and volumes and the variations of the SVs seen in Figure 5 are caused by the changes in thoracic pressure caused by the breathing reflex. As a spontaneous breath is triggered, the respiratory muscles increase the volume of the thoracic cavity, a pressure drop is generated and the RV venous return pressure increases. This causes an increase in SV in the RV and an increased inflow of blood into the lung. In parallel to this, the venous return to the LV decreases, leading to a decrease in the SV. During peak expiration the reverse occurs, the increased thoracic pressure creates a decrease in the venous return pressure and the RV stroke volume goes down, with the opposite happening in the LV. This is because the increased thoracic pressure generates an increase in pressure in the pulmonary capillaries and veins. The slightly increased thoracic pressure generates an overall decrease in venous return pressure and therefore an overall decrease in SV in both ventricles.

#### PPV

To simulate PPV, some modifications to the model parameters had to be made as these patients are usually sedated and paralysed. The effect of paralysis in muscles is to decrease its tonus; In our model tonus is positively correlated with V0 of the thorax and pleura and negatively correlated with their compliance. To replicate this, the maximum and minimum value of thorax *V*_0_ was set to 3200 and pleura *V*_0_ to 2800. The compliance of the pleura was also increased to 90 *mL/mmHg*. A ventilator was then connected to the mouth with a PEEP of 5mmHg (6.79*cmH*_2_*O*) and a breath amplitude of 7.3mmHg (10*cmH*_2_*O*). In Figure 6, the comparison between SB and PPV is presented.

**Figure 6.**
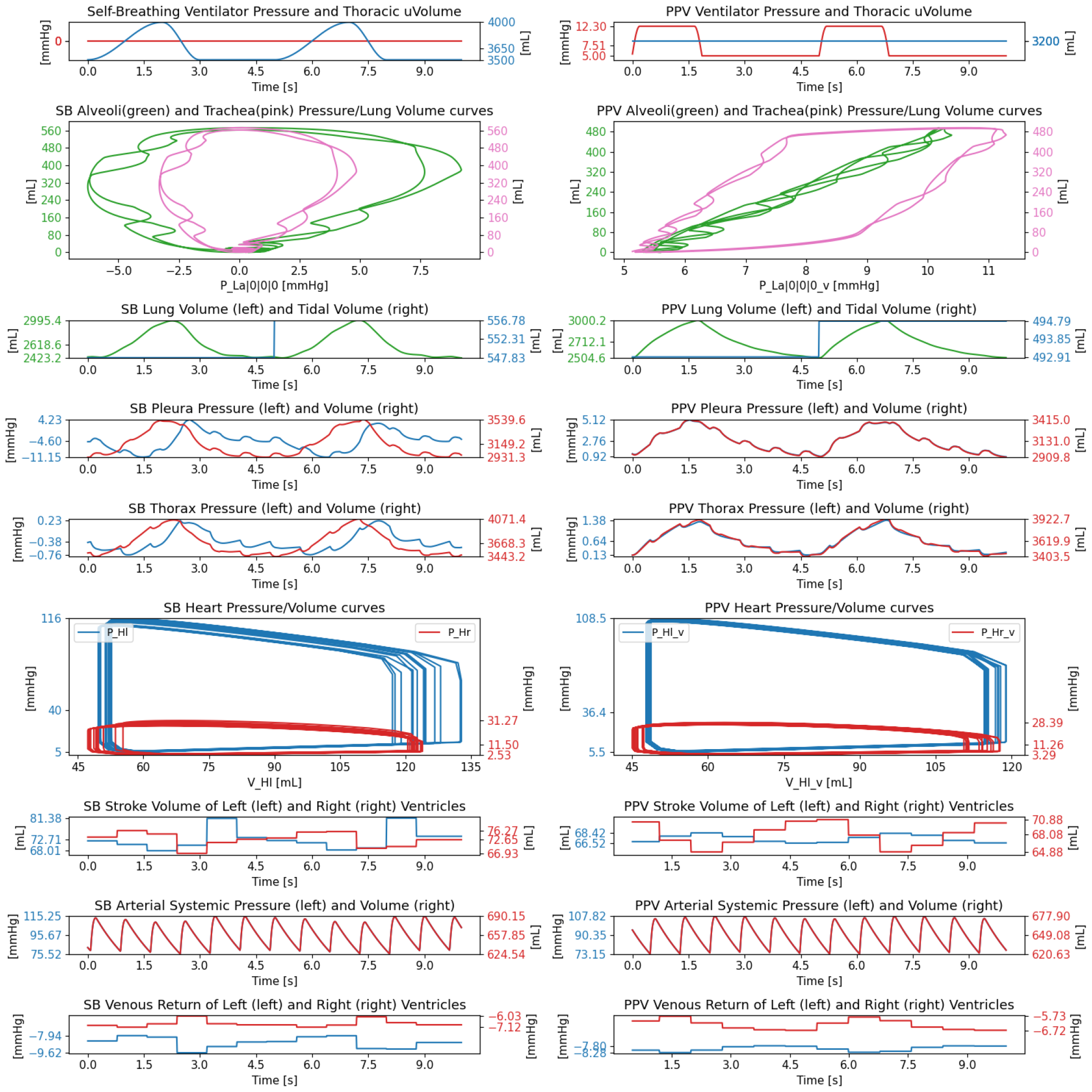
Comparison between self-breathing SB (left side) and positive pressure ventilation PPV over two breathing cycles. The *y−*axis for all plots displays the maximum, average and minimum values observed during the simulation.

In self-breathing mode, the average thoracic pressure was found to be -0.38 *mmHg*, with a minimum of -0.76 *mmHg* at the peak of inspiration PI and a maximum of 0.23 *mmHg* at the peak of expspiration PE along with a pleural pressure PlP ranging from -11.15 to 4.23 *mmHg*. This generated a Tidal Volume TV of 550 *mL* and a Functional residual capacity of 2423 *mL*. In PPV mode, the thoracic pressures varied between 0.13 to 1.38 *mmHg* with PlP ranging from 0.92 to 5.12 *mHg* and a TV of 493 *mL*. The average SV was also reduced from 72 *mL* during SB to 68 *mL* in PPV, resulting in a reduction of mean systemic BP of 5 *mmHg*.

In contrast to spontaneous breathing, during PPV the pressure in the thoracic cavity is highest at peak inspiration and therefore the SV is the lowest at this point and the highest during peak expiration. Another factor bringing the SV down during PPV is the increased blood pressure in the lung capillaries. Because the maximum elastance of the ventricles is constant, the RV does not generate enough force to maintain CO and this leads to a larger volume in the RV during diastole. Because of the reduced venous return to the LV during PPV, the blood volume in the LV also reduces leading to a lower systemic arterial blood pressure.

#### Heartbeat comparison

In the work of [37] the effect of the breathing reflex on the heart in healthy subjects is presented. Here the authors compare the Ejection Fraction (EF), Stroke Volume (SV) and End-Diastolic Volume (EDV) of both ventricles at peak inspiration and peak expiration. Table 9 presents the results of our model simulation alongside the values observed in [37]. The model appears to underestimate the changes of SV and EDV for the right ventricle but keeps well within the observed range for the expected drop in the left ventricle values. The literature also describes marginal changes in EF, which the model also seems to reproduce. During PPV, the system appears to homogenise the heartbeat during the breathing cycle, with fewer differences observed between peak inspiration/expiration. As expected these tendencies in PPV invert when compared to SB, with SV experiencing its minimum at peak inspiration and maximum at peak expiration.

**Table 9.** Changes in Stroke Volume SV, Ejection Fraction EF and end-Diastolic Volume SV in the Right Ventricle RV and Left Ventricle LV during the RC. The values presented are the percentage change from the average values during Peak Inspiration and Peak Expiration, The values in parenthesis are the distribution observed in [37]

|  | average<br>Self-Breathing | % change from average |  | average<br>ventilator | % change from average |  |
| --- | --- | --- | --- | --- | --- | --- |
|  |  | Peak Ins | Peak Exp |  | Peak Ins | Peak Exp |
| LV EF (%) | 59.2 | -1.82 (0) | -2.18 (0) | 58.6 | -0.84 | 1.02 |
| LV SV (mL) | 74.8 | -3.46 (-4 +-7) | -8.48 (+4 +-7) | 68.4 | -2.72 | 1.91 |
| LV EDV (mL) | 126.2 | -1.48 (-3) | -6.04 (+5 +-7) | 116.7 | -1.84 | 0.93 |
| RV EF (%) | 59.3 | 0.35 (0) | 2.83 (0) | 59.8 | 1.21 | -1.58 |
| RV SV (mL) | 72.6 | 0.55 (+10 +-5) | 3.96 | 68.5 | 2.91 | -1.46 |
| RV EDV (mL) | 122.5 | 0.14 (+9) | 1.11 (-9) | 114.6 | 1.68 | 0.07 |

### PEEP Challenge test

To perform the PEEP challenge, a ventilator is connected to the model with an initial PEEP of 0 *mmHg* and breath amplitude of 7.3 *mmHg* (10 *cmH*_2_*O*). Initially, 5 breaths were allowed to happen at this PEEP, followed by a progressive increase of 1*mmHg* in PEEP every other breath. The first breath after a PEEP step yields a TV that is artificially increased as PEEP increase contributes to the breath amplitude. The simulation was run until a maximum PEEP of 35 *mmHg* was reached. The lung was deflated by gradually lowering PEEP back to atmospheric pressure. The results for this test are presented in Figure 7.

**Figure 7.**
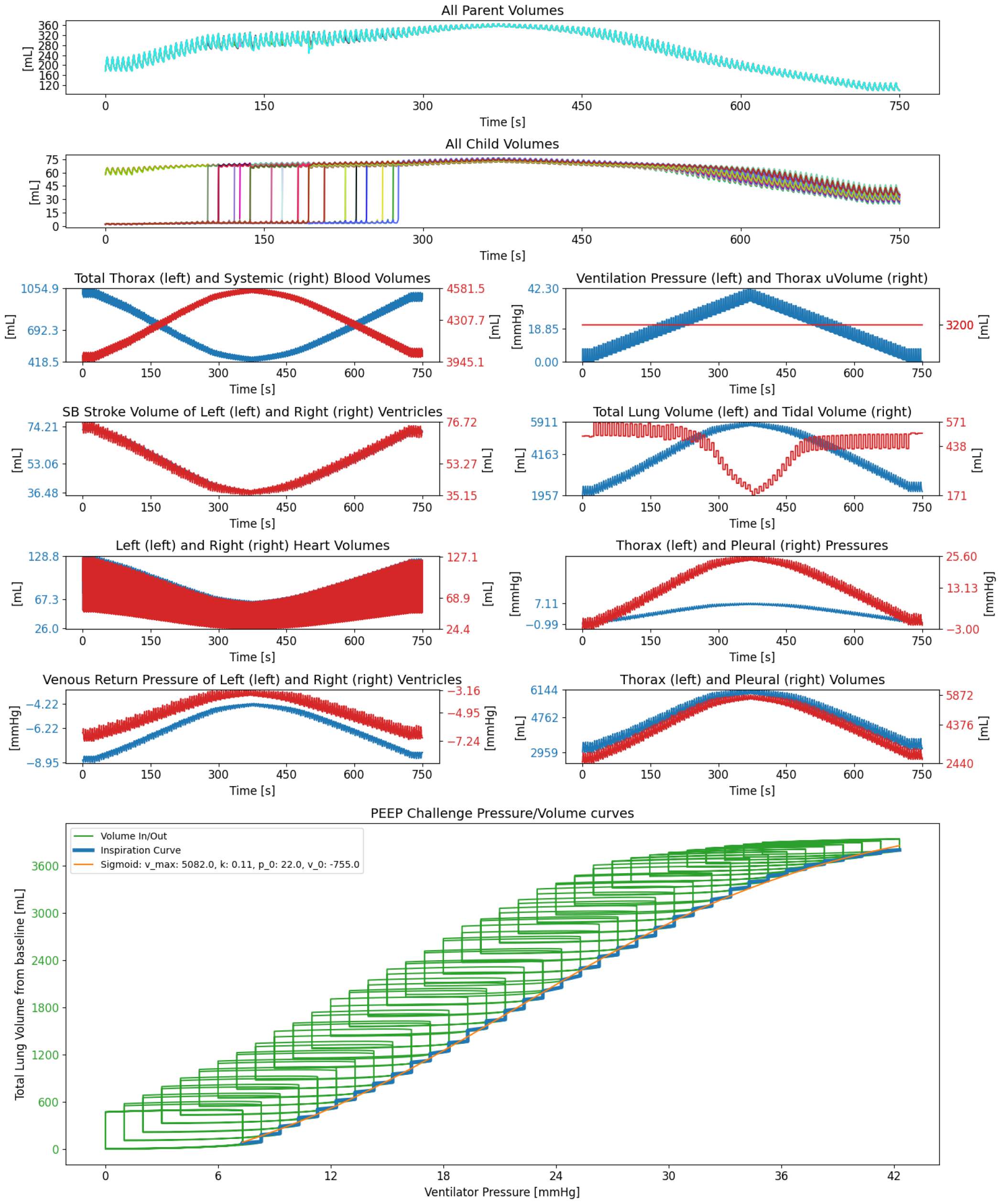
PEEP challenge test simulation displaying the systematic opening of the child alveoli as PEEP increases and its effect on the cardiovascular (left) and pulmonary (right) systems. In accordance with experimental data [38] [39] the system displays a sigmoidal relationship between volume and pressure (bottom)

In this conformation, the 8 parent alveoli are fully open initially, while the 32 child alveoli start fully closed and are systematically opened during the test as PEEP increases. This happens because the TOP of each alveolus is different, as illustrated in Figure 9.

As PEEP increases, the TV appears to remain relatively unchanged for the first few PEEP increments. A marked decrease in TV is observed for PEEP values around 23 *mmHg* (280 *s* into the simulation). This happens because at this pressure most alveoli are already opened and fully extended. Further volume increases will make the compliance drop and smaller changes in volume occur for the same pressure increase. In accordance with experimental data [38] [39], the developed model in this conformation is able to simulate a lung with a sigmoidal-like relationship between pressure and volume, as shown in the bottom plot of Figure 7. This relationship is highlighted by taking all the pressures and volumes at peak inspiration for all breaths of the inspiration phase of the test and fitting a sigmoidal curve to these points. According to the work of [40], the lung can start overstretching at 22 mmHg which, anecdotally, is the inflection point found by the curve fitting algorithm.

During the descending part of the PEEP test, the previously closed alveoli remained open, successfully simulating the opening effect that indicates why PEEP tests are sometimes prescribed to patients. Here, both the TV and FRC were also observed to increase slightly compared to their initial values; this is due to the total compliance of the lung being significantly higher and, consequently, the volume of air can now be redistributed through all the alveoli, and not only the parent ones.

The increase in PEEP generates an increase in thoracic and pleural pressures of 8 *mmHg* and *≃* 29*mmhg* respectively. The pressures in all cardiovascular compartments that are biased by these pressures will therefore also increase. This increase triggers an outflow of 600 *mL* of blood from the thoracic cavity into the systemic circulation. This shift of blood away from the lungs and the increasing external pressure felt by the lung capillaries make the volume in this compartment decrease. The increasing pressure in the system also decreases the systemic venous return pressure, reducing blood flow from the systemic circulation into the right ventricle. The stroke volume also decreases with increasing PEEP until it reaches 35 *mL* at maximum PEEP. This happens because the systolic elastance in the right ventricle does not generate enough pressure to overcome the pressure acting on the downstream vessels.

In Table 10 a comparison of the simulated lung volumes is presented alongside the values considered normal by the literature. Our simulated lung exhibits volumes close to, or within, the expected normal range.

**Table 10.** Volumes of the lung compared to the ones found in [36].

| Lung Volumes (mL) |  |  |
| --- | --- | --- |
| Name | Model Result | Normal Values [36] |
| Tidal Volume TV | 499-504 | 500 |
| Inspiratory reserve volume | 2500 | 3000 |
| Inspiratory capacity | 3000 | 3500 |
| Functional residual capacity | 2627 | 2300 |
| Total lung capacity | 5492 | 5800 |

### Deep Breath Test

A deep breath during spontaneous breathing is a comparable manoeuvre to the PEEP test, where the lung is filled with air to Total Lung Capacity (TLC). In this case, however, the respiratory muscles are responsible for the driving force of the breath instead of the ventilator. To test our model in this scenario a normal breath was allowed to happen, followed by a deep breath which is subsequently followed by another normal breath. The deep breath lasted 10 seconds (double the time of a normal breath) and was generated by increasing the value of *V*_0_ of the pleura and thorax compartments to 7150 *mL*. As can be seen in Figure 8, the alveoli dynamics are very similar to the PEEP test, where all parent alveoli are initially open while all child alveoli are closed. As *V*_0_ increases, the PlP decreases and the transpulmonary pressure increases past TOP. Consequently, the child alveoli start popping open in succession. At peak inspiration, the total lung volume was 5698 *mL* with a TV of 2983 *mL*, and the child alveoli remain open after deflation. On the cardiovascular side, the results are however very different. Instead of a net outflow of blood from the thoracic cavity, we see a net inflow of 200mL of blood during inspiration. The SV of the right ventricle increases during inspiration to a maximum of 89 *mL* and decreases during expiration to a minimum of 53 *mL* at peak expiration. The left ventricle follows an opposite trend ranging from a minimum of 52 *mL* during inspiration to a maximum of 93 *mL* at peak expiration. Despite a substantial increase in the volume of the thoracic cavity, the pulmonary circulation never ceased. The venous return into the right ventricle remains consistently negative and ventricular volumes are practically unchanged, other than a slight increase in left ventricular volume during expiration.

**Figure 8.**
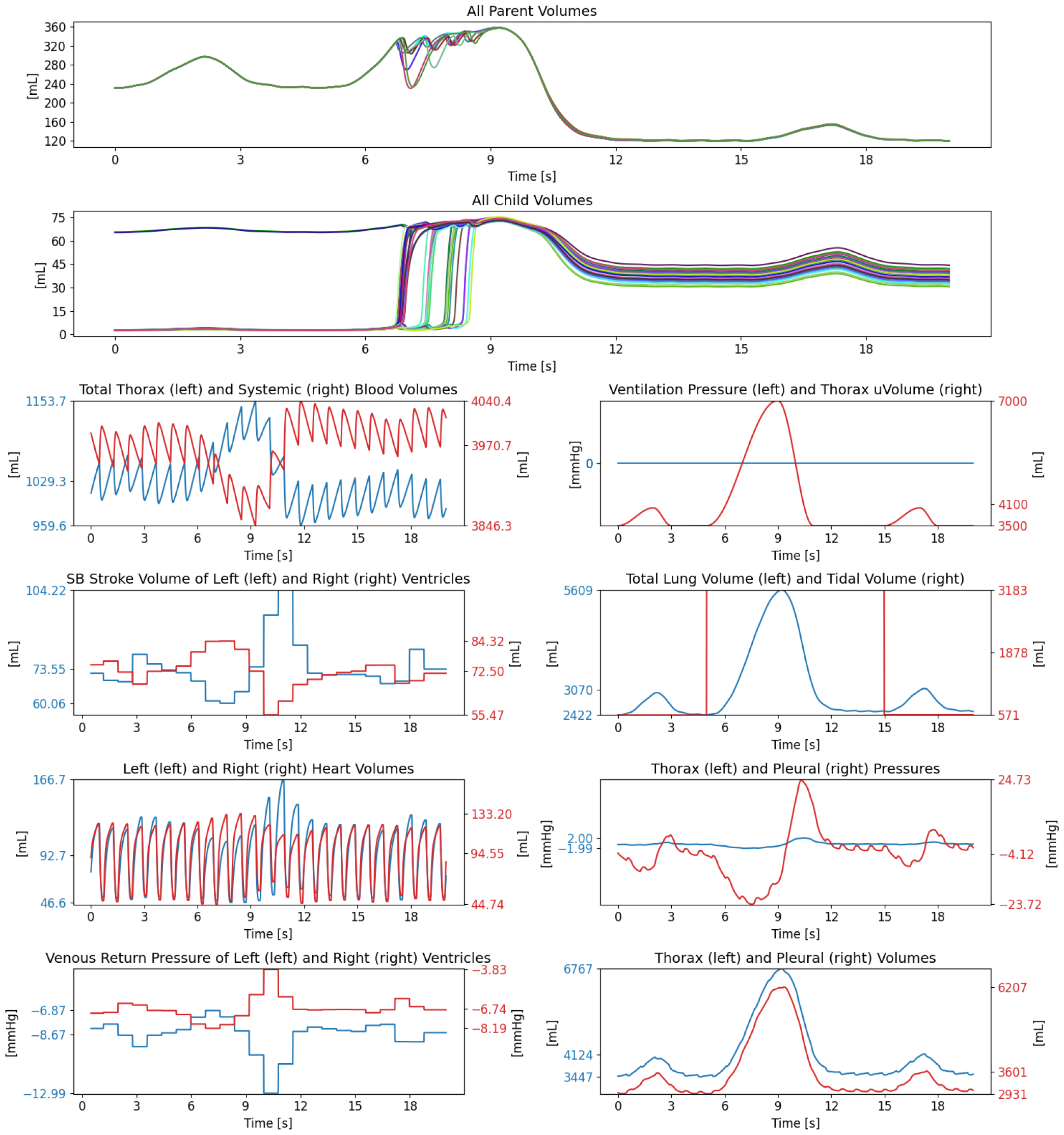
Deep breath Simulation displaying the volume redistribution of the parent alveoli as the child alveoli open due to the effect of a deep breath and its effect on the cardiovascular (left) and pulmonary (right) systems.

**Figure 9.**
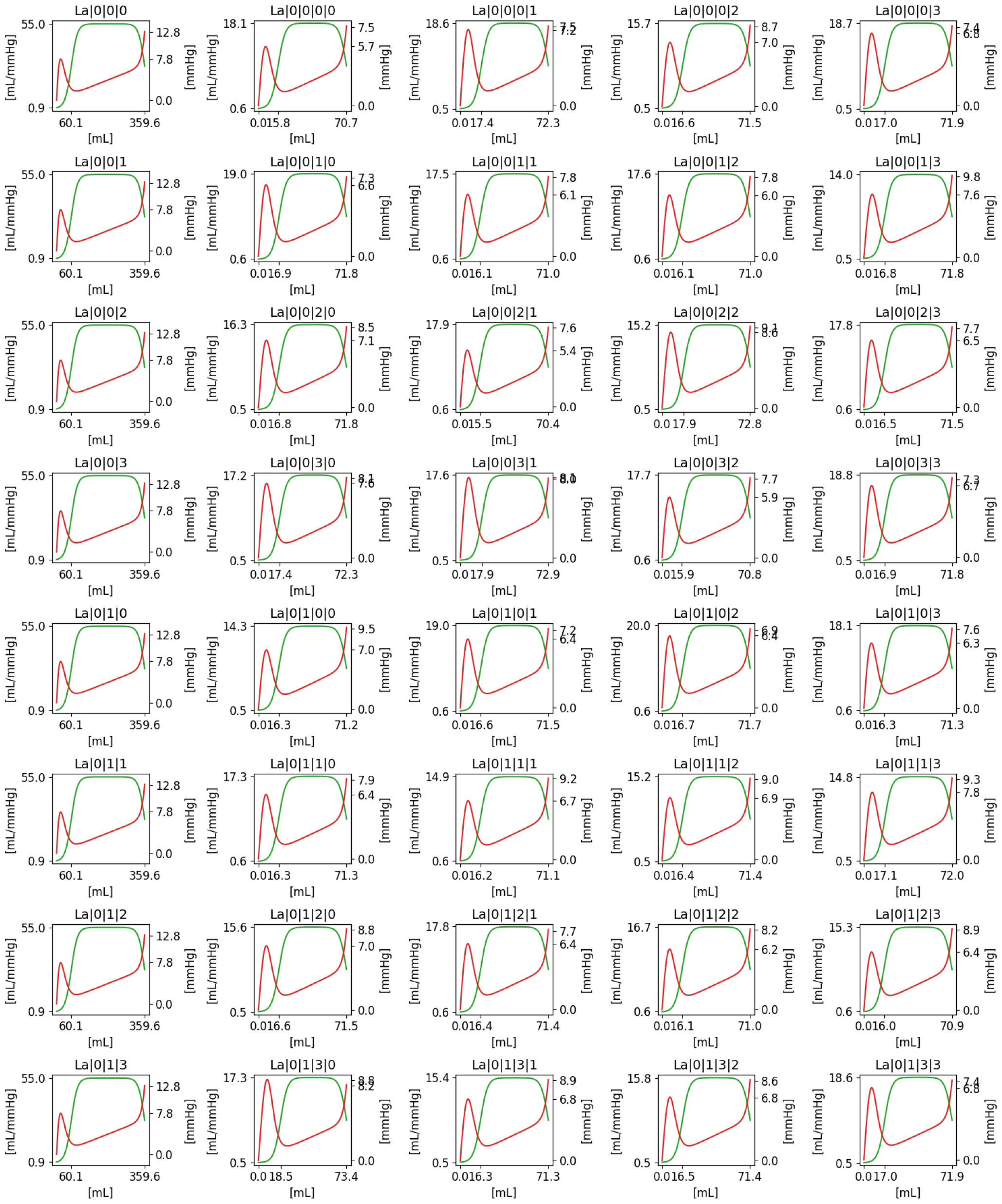
Compliance/Volume and Pressure/Volume curves for all 40 alveoli. the ticker in the y-axis is maximum and minimum compliance, minimum, TOP and maximum pressure and, for the x-axis, minimum, *V*_0*pop*_ and maximum volumes

## 4 DISCUSSION AND CONCLUSIONS

The approach outlined in this paper successfully captures the mechanical properties of the CP system, being capable of simulating most of the mechanical phenomena observed in [41] where the system can be observed to communicate changes in pressure and volume between pulmonary and cardiovascular compartments. This cross-talk between systems creates pressure gradients in the cardiovascular system resulting in shifts of blood volume between the pulmonary and the systemic circulation and changes in venous return pressure to the heart, leading to variations in stroke volume between heartbeats happening at different phases of the respiratory cycle. The proposed alveoli opening system is also observed to have an appropriate reaction to the PEEP challenge, where the lung is observed to open systematically with increasing PEEP generating the familiar sigmoidal pressure/volume profile and substantially affecting the cardiovascular system, where a marked decrease in CO is observed. In contrast, the reaction of the system to a deep breath was observed to have a much less detrimental effect in the cardiovascular function.

The modelling approach presented not only simplifies the understanding and interpretation of individual equations by allowing their independent analysis but also facilitates the creation of complex, highly interconnected models with extensive equation systems without requiring changes to the code base of the model. These features also allowed us to create a calibration strategy that can be used to generate models with different phenotypes enabling the study of different conditions and how they affect the system.

### Model limitations and the need for ANS

Despite the overall behaviour of the system being very close to what is seen in real patients, there are still some aspects where the model deviates from what is observed in clinical practice. Firstly, the model features a high-level representation of the physiological system with the whole cardiovascular CV system being summarised into very few compartments, in contrast to the millions of vessels and alveoli present in real humans. The absence of abdominal pressure, as well as the full accountability of inertial effects and blood rheology, the absence of ventricular interaction via the septum wall and pericardium, gravity and position all contribute, to a greater or lesser degree, to the real-world dynamics. The absence of these mechanism is mainly noticeable in the heart response to the effects of breathing where, despite the model displaying the correct overall dynamics, it underpredicts the variation of SV and EDV during a RC, as can be seen in Table 9.

Moreover, the model’s capability to simulate disease conditions or replicate physiological manoeuvres, such as exercise physiology or a Valsalva manoeuvre, is currently constrained by the absence of an autonomic nervous system (ANS). The adaptability of the human body to changes is pivotal in clinical scenarios, and the model’s limitation in incorporating these effects hinders the realism of certain tests, like the positive end-expiratory pressure (PEEP) test, where the absence of regulatory effects leads to a complete collapse of the pulmonary vessels. Incorporating ANS into the model would enhance its simulation capabilities, enabling a more accurate representation of the body’s responses to various stimuli, diseases, and medical procedures. A realistic approach to the development of an ANS would also require the addition of gas exchange and gas transport into the model, as one of the most important roles of ANS, in this context, is to ensure sufficient oxygenation to the vital organs and other tissues.

### Conclusions

The combined advantages of leveraging Python’s versatility and integration capabilities, coupled with the model’s configuration flexibility using JSON, facilitate the creation of user-friendly applications. This unique blend of technical strengths positions the model as an optimal choice for seamless integration and effective utilization within clinical environments. Additionally, this setup facilitates the creation of a distributed system, enabling the offloading of computational burdens to servers. Consequently, our proposed approach reduces the bedside requirements to a mere tablet and allows for the creation of web/mobile applications.

The object-oriented and modular approach used in the implementation of the system also allows for extensions to the model to be easily implemented. As proved by the use of the additional equations used to calibrate the model, all parameters (*R,C,V*_0_, *E*_*max*_, *E*_*min*_, *HC, RC*) used in this work can be easily converted into state variables, allowing most implementations of autonomic nervous system interactions found in the literature to be easily incorporated into our model and different regulation approaches to be tested. Allowing different model configurations and interactions to be configured rather than implemented.

The use of a first-principle and modular approach to the creation of the ODE system allows for the use of relatively simple equations that do not require extensive mathematical skills to comprehend, making the whole model easier to interpret.

#### Model applicability to clinical settings and as a research tool

One of the best potential uses for these types of mechanistic models is in patient-specific medical research. In this context, the model presented would constitute a skeleton where the parameters could be adjusted to represent the physiology of a particular patient. Finding the combination of parameters needed to represent the physiology of a specific patient will require more sophisticated parameterization methods; this will be the focus of future publications. A key part of the challenge to adopt these models in patient-specific studies is the nature of the data available to calibrate the models. In the spectrum of time series, routine clinical practice tends to only generate data, at a sufficient resolution for very specific variables in the model such as arterial systemic BP, HR, RR, TV and central venous pressure. If the optimisation algorithms attempt to blindly find a combination of model parameters that can generate the same trends on the variables recorded from the patient, the likelihood of different combinations of parameters with very different physiological interpretations emerging is very significant. It is therefore important to test the capabilities of these algorithms to disambiguate between different parameter regimes. The framework for model creation presented in this work is ideal for this type of application as it provides the structural flexibility needed to test different scenarios, approaches and effects.

## APPENDIX

## ACKNOWLEDGMENTS

The authors are grateful for the funding and support of the EPSRC-funded CHIMERA Maths in Healthcare Hub (EP/T017791/1), the Wellcome/EPSRC Centre for Interventional and Surgical Sciences (WEISS) (203145/A/16/Z), the Department of Mechanical Engineering and Department of Mathematics at UCL.

## GLOSSARY

ARDS: Acute Respiratory Syndrome. 2
BP: Blood Pressure. 11–13, 16
CO: Cardiac Output. 2, 11, 14, 15
CP: cardiopulmonary. 2, 5, 8, 11, 15
CV: cardiovascular. 2, 5, 10, 11, 15
EDV: end-diastolic volume. 14, 16
EF: ejection fraction. 14
FRC: Functional Residual Capacity. 11, 14
LV: Left Ventricle. 13, 14
PE: Peak Expiration. 13
PEEP: positive-end-expiratory pressure. 2, 8, 13–15
PI: Peak Inspiration. 13
PlP: Pleural Pressure. 13, 15
PPV: Positive Pressure Ventilation. 2, 13, 14, 17
PS: pulmonary system. 2, 5, 11
RC: Respiratory Cycle. 8, 11, 14, 16
RV: Right Ventricle. 13, 14
SB: spontaneous breathing. 2, 13, 14, 17
SV: Stroke Volume. 11–16
TLC: Total Lung Capacity. 15
TOP: Threshold Opening Pressure. 2, 6, 11, 14, 15
TV: Tidal Volume. 13–16

## Footnotes

1 calculated using the knowledge of the duration of a RC and the Inspiration/Expiration (I:E) ratio defined on the ventilator

## Notes

### Competing Interest Statement

The authors have declared no competing interest.

### Summary of Updates

Acknowledgements were incomplete and are ow updated with: The authors are grateful for the funding and support of the EPSRC-funded CHIMERA Maths in Healthcare Hub (EP/T017791/1), the Wellcome/EPSRC Centre for Interventional and Surgical Sciences (WEISS) (203145/A/16/Z), the Department of Mechani- cal Engineering and Department of Mathematics at UCL.

